# An iMSC-Based iPSC Model for Osteogenesis Imperfecta: A Platform for Disease Modeling and Drug Screening

**DOI:** 10.1101/2025.06.11.659155

**Authors:** Ashis Kumar, Vignesh Kumar, Agnes Selina, Vrisha Madhuri, Vasanth Thamodaran

**Affiliations:** Centre for Stem Cell Research, a unit of inStem, Bengaluru, Christian Medical College, Vellore, Tamilnadu - 632002; Sree Chitra Tirunal Institute for Medical Sciences & Technology, Thiruvananthapuram - 695011, Kerala, India; Department of Paediatric Orthopedics, Amara Hospital, Tirupati, Andhra Pradesh - 517520, India; Tata Institute for Genetics and Society, inStem, Bengaluru, Karnataka - 560065; Academy of Scientific and Innovative Research (AcSIR), Ghaziabad, Uttar Pradesh - 201002

## Abstract

**Introduction:** Osteogenesis Imperfecta (OI) is a rare genetic disorder of connective tissue, primarily caused by mutations in the COL1A1 or COL1A2 genes. Research is hampered by the limitations of primary patient-derived cells, which are obtained through invasive methods and have finite proliferative capacity and high variability. While induced pluripotent stem cell (iPSC) models exist, direct differentiation to osteoblasts is often inefficient and may not fully replicate disease characteristics. Our study aimed to develop and characterize a robust iPSC-derived mesenchymal stem cell (iMSC) model of OI using a simple two-step differentiation protocol to serve as a platform for disease modeling and drug screening.

**Methods:** Bone marrow MSCs (BMMSCs) were isolated from two OI patients with heterozygous missense mutations, in COL1A1 (c.2299G >A) and COL1A2 (c.982G>A). The patient BMMSCs were reprogrammed into iPSCs using an integration-free Sendai virus. The resulting OI-iPSCs were characterized for pluripotency via immunofluorescence and RT-PCR, trilineage differentiation potential, and karyotype analysis. A two-step protocol was used to differentiate the OI-iPSCs into OI-iMSCs via mesodermal lineage. The OI-iMSCs were then characterized for morphology, immunophenotype by flow cytometry, and trilineage differentiation capacity. Functional osteogenic differentiation was assessed by Alizarin Red S staining and RT-PCR analysis of key osteogenic genes. Sanger sequencing confirmed the retention of patient-specific mutations in the OI-iMSCs.

**Results:** Patient-derived BMMSCs were successfully differentiated into adipocytes and chondrocytes but showed impaired osteogenic potential. The reprogramming process successfully generated stable OI-iPSC lines that expressed key pluripotency markers, demonstrated trilineage potential, and maintained a normal karyotype. Critically, while direct osteogenic differentiation of these iPSCs failed to recapitulate the primary cell phenotype, the two-step differentiation protocol successfully produced a homogeneous population of iMSCs. These OI-iMSCs displayed characteristics of MSC surface markers (CD73+, CD90+, CD105+) and recapitulated the disease phenotype seen in the original patient BMMSCs. Specifically, COL1A1-iMSCs exhibited cellular rolling and detachment, while COL1A2-iMSCs showed poor mineralization during osteogenic differentiation. Both OI-iMSC lines showed significantly decreased calcium deposition and downregulation of key osteogenic genes (RUNX2, ALP, COL1) compared to controls.

**Conclusion:** Our study successfully established an iMSC-based cellular model of OI that recapitulates patient-specific disease phenotypes, including impaired osteogenic differentiation. The two-stage differentiation of iPSCs to iMSCs proved more reliable than direct osteogenic differentiation for modeling the disease. iMSC model circumvents the limitations of primary cells by providing a scalable and homogeneous source of patient-specific cells. Our platform offers a valuable and robust tool for investigating OI pathophysiology and for high-throughput screening of potential therapeutic molecules, advancing efforts toward personalized therapies

## Introduction

Osteogenesis Imperfecta (OI) is a rare, genetically heterogeneous connective tissue disorder primarily characterized by low bone mass, bone fragility, and skeletal dysplasia. The majority of OI cases, approximately 85%, are caused by pathogenic mutations in the COL1A1 or COL1A2 genes, which encode the α1(I) and α2(I) chains of type I collagen, a critical component of the bone matrix ^1,2^. These mutations disrupt the structural integrity of collagen, leading to brittle bones and impaired osteogenic differentiation ^2^. Osteoblasts are bone-forming cells that produce type I collagen and play a central role in the manifestation of OI. These cells offer valuable information to understand the pathophysiology of OI ^3^. Primary osteoblasts, essential for in vitro bone tissue studies, are typically obtained from bone biopsies or orthopedic surgeries. However, these methods are associated with considerable patient morbidity, including procedural discomfort and surgical risks ^4,5^. In addition, the utilization of primary MSCs in the investigation of rare bone disorders is constrained by several limitations. Specifically, primary MSCs exhibit a finite proliferative capacity and diminished differentiation potential during extended culture periods. Significant inter-individual variability in donor cell proliferation and differentiation potential further compounds these limitations ^6^. These limitations can be overcome by using induced pluripotent stem cell based (iPSC) disease models. iPSCs can be efficiently differentiated to MSCs, and the iMSCs are found to have better proliferative capacity with tri-lineage differentiation potentials comparable to adult MSCs^7^.

iPSC based cellular disease models of OI have been shown to recapitulate some of the disease phenotypes. Osteoblasts derived from OI patient iPSCs have shown defects in mineralization and collagen synthesis ^8^. Direct differentiation of iPSCs to osteoblasts often leads to incomplete or poor osteogenic differentiation and low yield of osteogenic cells ^9^. Induced mesenchymal stem cells (iMSCs) derived from iPSCs can offer a robust platform for studying the pathophysiology of OI and identifying potential therapeutic interventions^8,10,11^. iMSCs derived from iPSCs through embryoid body formation and neural crest lineage specification exhibit MSC-specific marker expression and trilineage differentiation potential into osteoblasts, adipocytes, and chondrocytes. However, these methods are limited by low efficiency and cellular heterogeneity during embryoid body formation^12^. During embryonic development, MSCs differentiated from lateral plate mesoderm significantly contribute to the formation of skeletal muscle and connective tissue^13^. Hence, differentiating iPSCs via mesodermal lineage to MSCs would be ideal for studying connective tissue disorders. In this study, we aimed to establish and characterize an iMSC-based cellular disease model of OI harboring genetic variants in COL1A1 and COL1A2 genes. This model, developed via a simple two step differentiation protocol involving mesoderm specification followed by iMSC derivation can serve as a robust platform for both studying OI pathophysiology and screening potential therapeutic drug molecules.

## Materials and methods

### Sample collection

Bone marrow aspirates were obtained from osteogenesis imperfecta patients undergoing elective surgery for deformity correction. Patients were screened and identified based on the genetic analysis. Informed written consent was obtained in accordance with the Declaration of Helsinki. The study was approved by the institutional ethics committee, Christian Medical College, Vellore.

### Clinical History and Genetic Phenotype

Patients were categorized based on severity of symptoms using the Sillence classification system. The first patient with a COL1A1 variant presented with type IV OI, and the second patient with a COL1A2 variant presented with type III OI. Both the patients presented with multiple fractures (>5), blue sclera, wormian bone, lower or upper limb deformity, and low bone mineral density. Clinical exosome sequencing revealed heterozygous single base exchange in COL1A1:c.2299G>A and COL1A2:c.982G>A, resulting in missense mutations p.(Gly767Ser) and p.(Gly328Ser) causing severe phenotype.

### Isolation and culture of bone marrow MSC

Mesenchymal stromal cells were isolated by standard protocol using the ficoll plague density gradient method ^14^. Briefly, 2 mL of bone marrow aspirate was diluted to 10 mL with phosphate buffered saline and layered gently on Ficoll at a 1:3 ratio and centrifuged at 1500 g for 30 mins at 4°C. The buffy coat containing the mononuclear cell population was collected and washed twice with phosphate-buffered saline (PBS). The cell pellet was resuspended in Dulbecco’s Modified Eagles Medium-low glucose (DMEM-LG) + 20% fetal bovine serum + antibiotics and seeded at a cell density of 10,000 cells/sq. cm. Flasks were maintained at 37°C in a humidified environment with 5% CO2. After 4 days, the unattached cells were removed, and the medium was replaced. The medium was changed every 3–4 days subsequently. At 70 to 80% confluence, the cells were detached using 0.05% trypsin and passaged at a density of 4 × 10^3^ cells/cm^2^ in culture flasks. Starting from the first passage, the cells were trypsinized at 80% confluence and replated at a density of 6000 cells/cm^2^. At passage 3, the MSCs were subjected to characterization by immunophenotyping and trilineage differentiation.

### Generation of iPSCs using integration-free virus

Approvals for deriving iPSCs were obtained from both human ethics and stem cell committee at the Tata Institute for Genetics and Society. To derive OI-iPSCs, BM-MSCs from COL1A1 and COL1A2 OI patients at passage 3 were reprogrammed using the Cytotune™ iPSC 2.0 Sendai Reprogramming Kit (Life Technologies). Briefly, 1×10^5^ BM-MSCs were seeded per 6-well plate and incubated at 37°C and 5% CO2. On day 2, the cells were transduced with the Sendai virus vector carrying the Yamanaka factor (OCT3/4, SOX2, KLF4, and c-MYC) in MSC growth medium at an MOI of 5 according to the manufacturer guide. Fresh MSC medium was replaced after 24 hours of transduction and cultured for 6 days in MSC medium. On day 7, the cells were trypsinized, and 5 × 10^5^ cells were plated on a 35 mm dish coated with Geltrex and cultured with MSC growth medium. After overnight culture in MSC growth media, the medium was replaced every alternate day with mTESR plus media (Stem Cell Technology). The iPSC colonies were picked for expansion starting from day 16-20 of reprogramming. Characterization of iPSCs was done at passage 10-15 after expansion.

### Characterization of iPSC for pluripotent markers

#### Immunofluorescence of iPSC markers

The two iPSC cell lines derived from COL1A1 and COL1A2 MSCs were fixed with 4% paraformaldehyde and subjected to immunofluorescence with pluripotent markers SSEA4, TRA1-60, TRA1-81, OCT4, SOX2, and NANOG (Cell Signaling Technology, MA, USA).

#### Trilineage differentiation assay and immunofluorescence

The two iPSC cell lines derived from COL1A1 and COL1A2 MSCs were subjected to trilineage differentiation into ectoderm, endoderm, and mesoderm lineages using the STEMdiff trilineage differentiation kit (Stem Cell Technologies, Canada). Briefly, single-cell suspensions of iPSC cell lines were seeded in Geltrex-coated culture plates with 10 µM of Y-27632 in mTeSR+ medium on day 0. The medium was replaced with appropriate differentiation medium (i.e., ectoderm, endoderm, and mesoderm differentiating medium) after 24 hours of seeding and replaced with fresh medium every alternate day until day 5 for mesoderm and endoderm lineage and until day 7 for ectoderm lineage. At the end of differentiation, the cells were fixed using 2% paraformaldehyde for 10 mins, followed by immunostaining with anti-PAX6 (ectoderm), anti-Brachyury (mesoderm), and anti-SOX17 (endoderm) antibodies to confirm the presence of lineage-specific markers.

#### Karyotyping

iPSC and iMSC were cultured in a 25 cm^2^ flask for 72 hours and treated with 10 µg/µL of colcemid (Sigma-Aldrich, NJ, USA) for 30 minutes, followed by hypotonic shock with 60 mM potassium chloride and fixed with methanol. Chromosomes of at least 10 to 20 metaphases were counted under an optical microscope. Karyotyping was performed at passage 10 for iPSCs and passage 5 for iMSCs.

#### Differentiation of iPSC to iMSC

In the phase-I of differentiation, the two iPSC cell lines derived from COL1A1 and COL1A2 patients were expanded in Geltrex-coated culture plates with mTeSR plus media. At 70% confluency, the media was replaced with STEMdiff™ mesenchymal progenitor induction medium (StemCell Technology, Canada) and cultured for four days to differentiate iPSCs to early mesodermal progenitor cells. In the second phase, the medium was replaced with animal component-free mesenchymal progenitor cell differentiation medium and cultured for 2 days to generate early mesenchymal progenitor cells. The cells were passaged and expanded on precoated culture plates with alpha minimum essential medium supplemented with 10% fetal bovine serum and 5 ng/mL fibroblast growth factor (Peprotech, NJ, USA) to generate functional induced mesenchymal stem cells (iMSCs). At 70 to 80% confluence, the cells were detached using 0.05% trypsin and passaged at a density of 4 × 10^3^ cells/cm^2^ in culture flasks. Starting from the first passage, the cells were trypsinized at 80% confluence and replated at a density of 6000 cells/cm^2^. At passage 3, the iMSCs were subjected to characterization by immunophenotyping and trilineage differentiation.

### Characterisation of OI-iMSCs

#### Immunophenotyping of OI-iPSCs and OI-iMSCs

Culture-expanded iPSCs and iMSCs were subjected to flow cytometric analysis of the MSC surface markers and iPSC markers. The detached cells were washed and resuspended in phosphate-buffered saline. To identify surface receptors, the cells were stained with fluorescein-conjugated antibodies against CD73, CD90, CD105, CD31, CD45, HLA-DRII (BD Bioscience, NJ, USA), CD56, CXCR4, CD81, CD24, CD117, TRA-1-60, and SSEA4 and incubated for 15 min at room temperature in the dark. The cells were washed and resuspended in DPBS and then acquired using a flow cytometer (CytoFLEX-LX, Beckman Coulter, CA, USA).

#### Trilineage differentiation of mesenchymal stem cell

To evaluate the multipotency of the OI-iMSCs, their differentiation into adipogenic, chondrogenic, and osteogenic lineages was assessed following a previously established protocol (REF). Adipogenic potential was assessed after two weeks of induction in adipogenic medium (containing Dulbecco’s modified Eagle’s medium-high glucose (DMEM-HG), 10% FBS, 0.5 mM 1-methyl-3-isobutylxanthine, 10 μg/ml insulin, 0.2 μM indomethacin, and 1.0 μM dexamethasone; Sigma, MO, USA); successful differentiation was confirmed by staining for lipid accumulation using Oil Red O staining. Chondrogenic differentiation was induced for 21 days by culturing cells in Lonza chondrogenic medium supplemented with insulin-transferrin-sodium selenite media supplement (Sigma), 5.33 μg/ml linoleic acid, 1.25 mg/ml BSA, 0.17 mM ascorbate-2-phosphate, 0.1 μM dexamethasone (Sigma), 1 mM sodium pyruvate (Fluka AG, Buchs, SC, USA), 0.35 mM proline, and 10 ng/ml TGF-β3 (R&D Systems, MN, USA); differentiation was confirmed by Safranin O staining, which detects the presence of glycosaminoglycans. Finally, osteogenic differentiation was induced over 21 days by culturing cells in DMEM-LG medium supplemented with 0.05 U/ml penicillin, 0.05 μg/ml streptomycin (Invitrogen), 10% FBS, 0.05 mM ascorbic acid, 10 mM glycerophosphate, and 0.1 μM dexamethasone. The presence of mineralized matrix deposition, characteristic of bone formation, was assessed by staining calcium nodules with Alizarin Red S; this staining was subsequently quantified by eluting the dye with 10% cetylpyridinium chloride and measuring the absorbance at 405 nm.

#### Sequencing analysis

Genomic DNA was extracted from the patient MSCs and OI-iMSCs using the Qigen Blood DNA Kit (Qigen). Primers were designed to amplify the mutant region (primer sequences are described in Table 1). Amplicons were purified and sequenced in both directions using the BigDye Terminator Kit using an ABI 3730 Genetic Analyzer (Applied Biosystems).

**Table 1:**
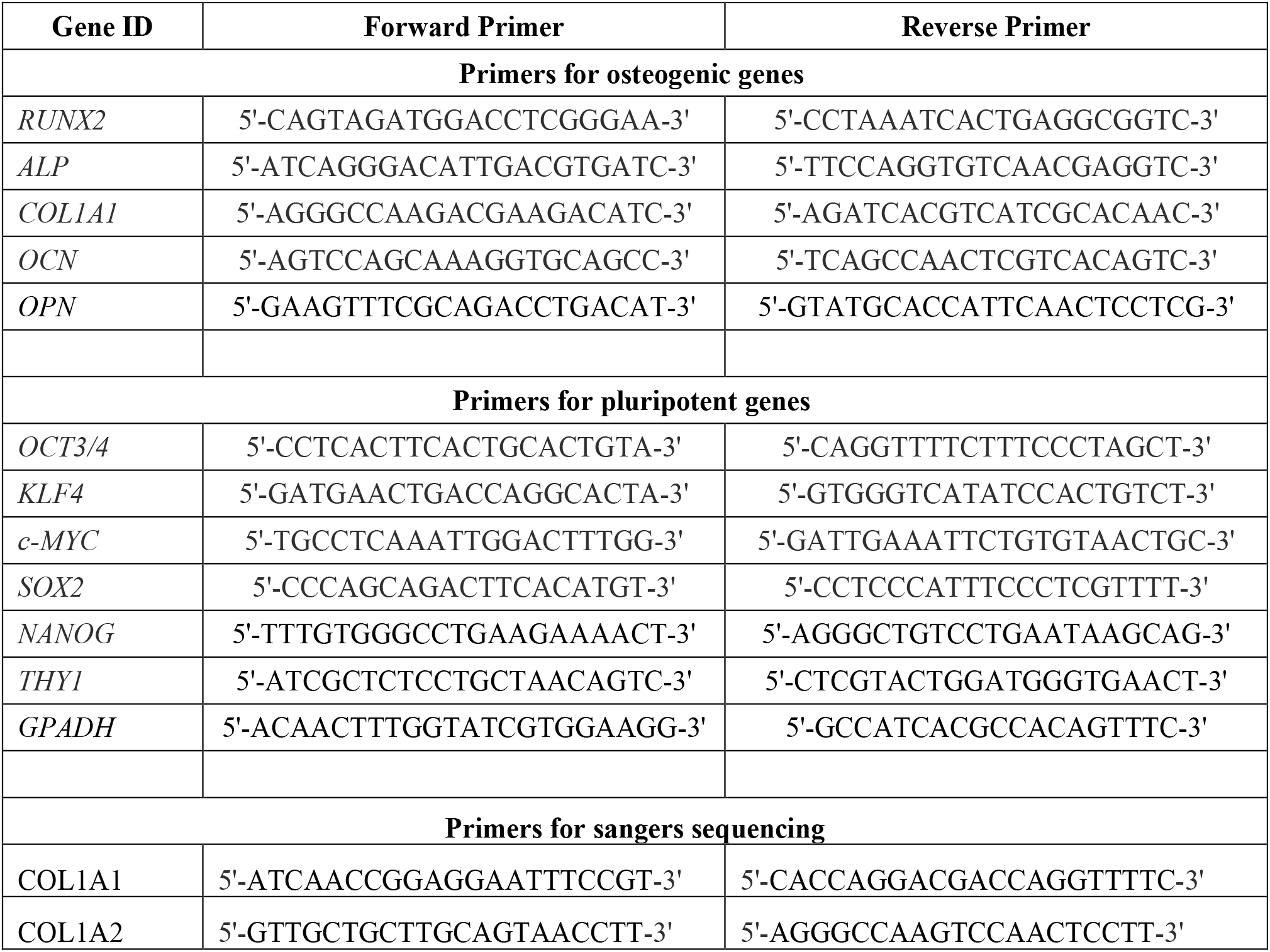
List of Primers.

#### Gene expression analysis

To assess the pluripotent gene expression of iPSCs, the total RNA of iPSCs from passage 10 was isolated and reverse transcribed using a PrimeScript First Strand cDNA Synthesis Kit (Takara, Japan). Amplification was performed using TB Green Premix Ex Taq II master mix (Takara) and the QuantStudio Real-Time PCR system (Thermo Fisher Scientific). According to the ISCT guidelines, primers were designed for 5 different pluripotent genes (primers listed in the table 1).

To assess the osteogenic differentiation, total RNA from day 7, day 14, and day 21 was isolated and subjected to real-time PCR. (The primers used are listed in the table 1.). The relative gene expression of the target genes was normalized to that of GAPDH.

## Results

### Characterisation of patient derived bone marrow MSCs

Mesenchymal stromal cells were isolated and cultured from bone marrow aspirated from two patients with mutations in the COL1A1:c.2299G>A and COL1A2:c.982G>A genes. Mononuclear cells were isolated using Ficoll-plaque, seeded at a density of 1 × 10^5^ cells/cm^2^, and cultured in **α** -MEM supplemented with 20% FBS and antibiotics. Starting from day 6-8, spindle-shaped fibroblast-like cells were observed (Fig. 1A). Trilineage differentiation potential was confirmed by adipocyte, chondrocyte, and osteoblast differentiation (Fig 1B). OI-BMMSCs with mutations in COL1A1 and COL1A2 genes demonstrated the accumulation of proteoglycans and glycosaminoglycans by safranin O staining, and the adipogenic phenotype demonstrated lipid vacuoles by oil red O staining. However, COL1A1-OI BMMSCs showed impaired osteogenic differentiation; the cells started to roll and fold during the osteogenic differentiation, while the COL1A2-OI BMMSCs differentiated to osteoblasts but showed poor mineralization with alizarin red staining (Fig 1B). To confirm the MSC phenotype, passage 3 cells were subjected to immunophenotyping. OI-BMMSCs (COL1A1 and COL1A2) showed more than 98% expression of positive markers CD90, CD105, CD73, and CD44 and expressed <1% of the negative markers CD34, CD45, and HLA-DR (Fig 1C). Gene expression analysis of key chondrogenic and adipogenic genes showed upregulation of SOX2, ACAN, PPARy, and FABP4, while the osteogenic genes showed downregulation of RUNX2, ALP, and OCN compared to healthy BMMSCs (p < 0.05, p < 0.005, p < 0.0001) (Fig 1D). Indicating a dysregulation in differentiation towards the osteogenic lineage.

**Figure 1.**
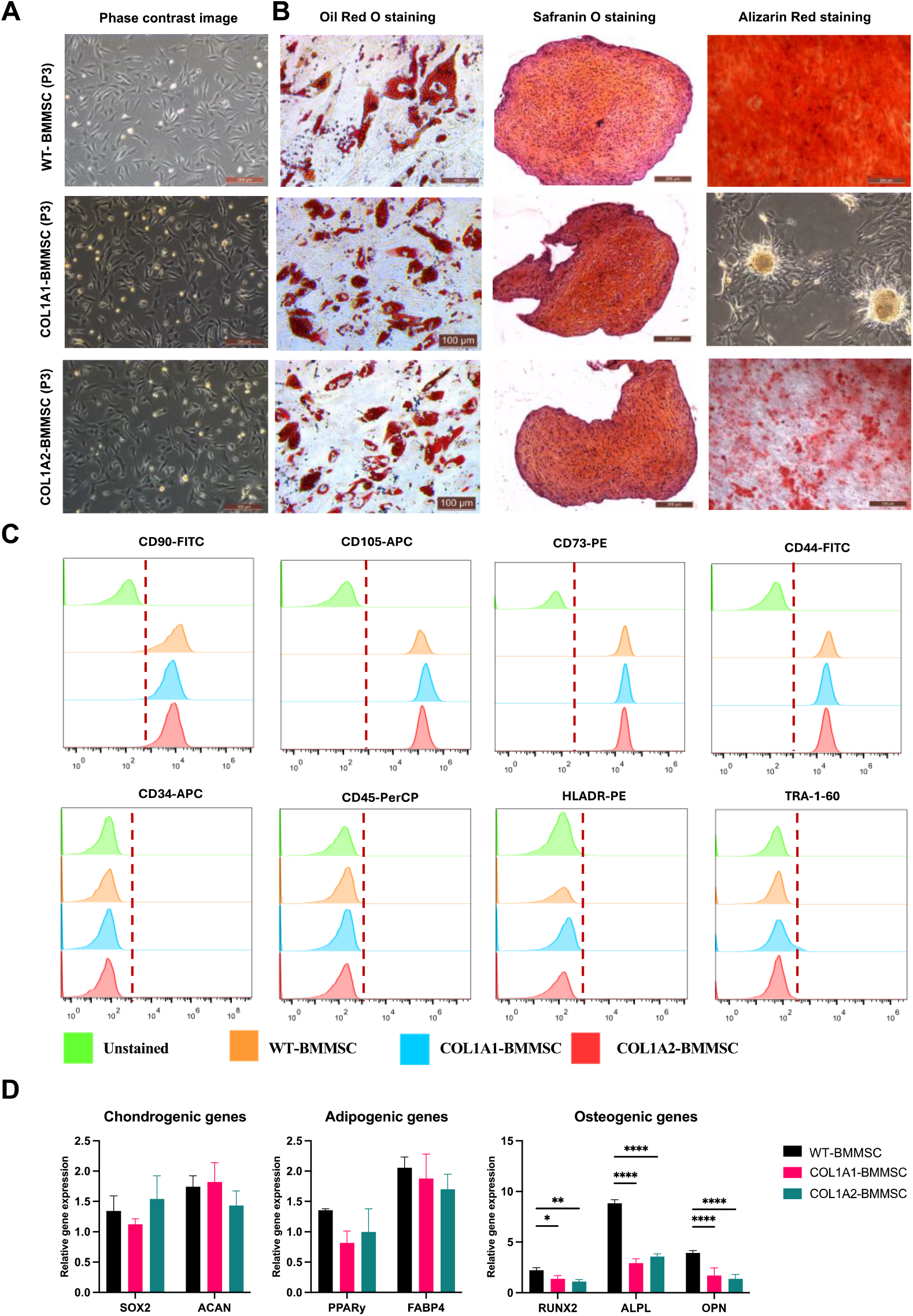
Characterization of patient-derived bone marrow mesenchymal stromal cells (BMMSCs). (A) Phase-contrast microscopy image shows the typical spindle-shaped, fibroblast-like morphology of OI-BMMSCs. (B) Trilineage differentiation potential demonstrated by oil red O staining reveals lipid vacuole accumulation (adipogenesis), Safranin O staining shows expression of glycosaminoglycan (chondrogenesis), and Alizarin Red staining demonstrates calcium deposition (osteogenesis). (C) Flow cytometric analysis confirms the expression of positive MSC surface markers (CD90, CD105, CD73, CD44) and lack of expression of negative hematopoietic and endothelial markers (CD34, CD45, HLA-DR). (D) Gene expression analysis following trilineage differentiation shows the expression of key chondrogenic (SOX2, ACAN) and adipogenic (PPARγ, FABP4) genes was confirmed/upregulated. Osteogenic genes (RUNX2, ALPL, OPN) were significantly downregulated in OI-BMMSCs compared to healthy control BMMSCs at day 21 (p < 0.05, *p < 0.001, ***p < 0.0001)

### Generation and characterization of pluripotent stem cells

Unlike primary MSCs, iPSC derived MSCs display better growth potential ^15^. In addition, the iPSCs ability to be maintained indefinitely offers an advantage of expanding them in large scale and storing the cells for any future disease modeling and drug screening studies. Given these advantages, iPSCs were generated from the OI-BMMSCs (P3) isolated from the two osteogenesis imperfecta patients with COL1A1:c.2299G>A and COL1A2:c.982G>A gene variants. Three to six clones were chosen and expanded for further characterization. All clones showed iPSC morphology, smooth-edged colonies with a scanty cytoplasm-to-nucleus ratio (Fig. 2A). The presence of stem cell markers such as *SOX2, OCT4, NANOG, SSEA4, TRA-1-60*, TRA-1-81, Lin28 and DMNT3B were evaluated by immunostaining and RT-PCR (Fig. 2B and C). Further, the ability of the cell lines to differentiate into ectoderm (PAX6), mesoderm (T-Brachury), and endoderm (SOX17) lineages demonstrated the successful reprogramming into pluripotent stem cells (Fig 2D). Karyotyping analysis revealed a normal karyogram, and no evidence of mycoplasma was observed (Fig. 2E and F). Our results demonstrate similar expression of pluripotent and trilineage differentiation in both OI-MSC-iPSC and hfMSC-iPSC (supplementary fig. 1A-E)

**Figure 2.**
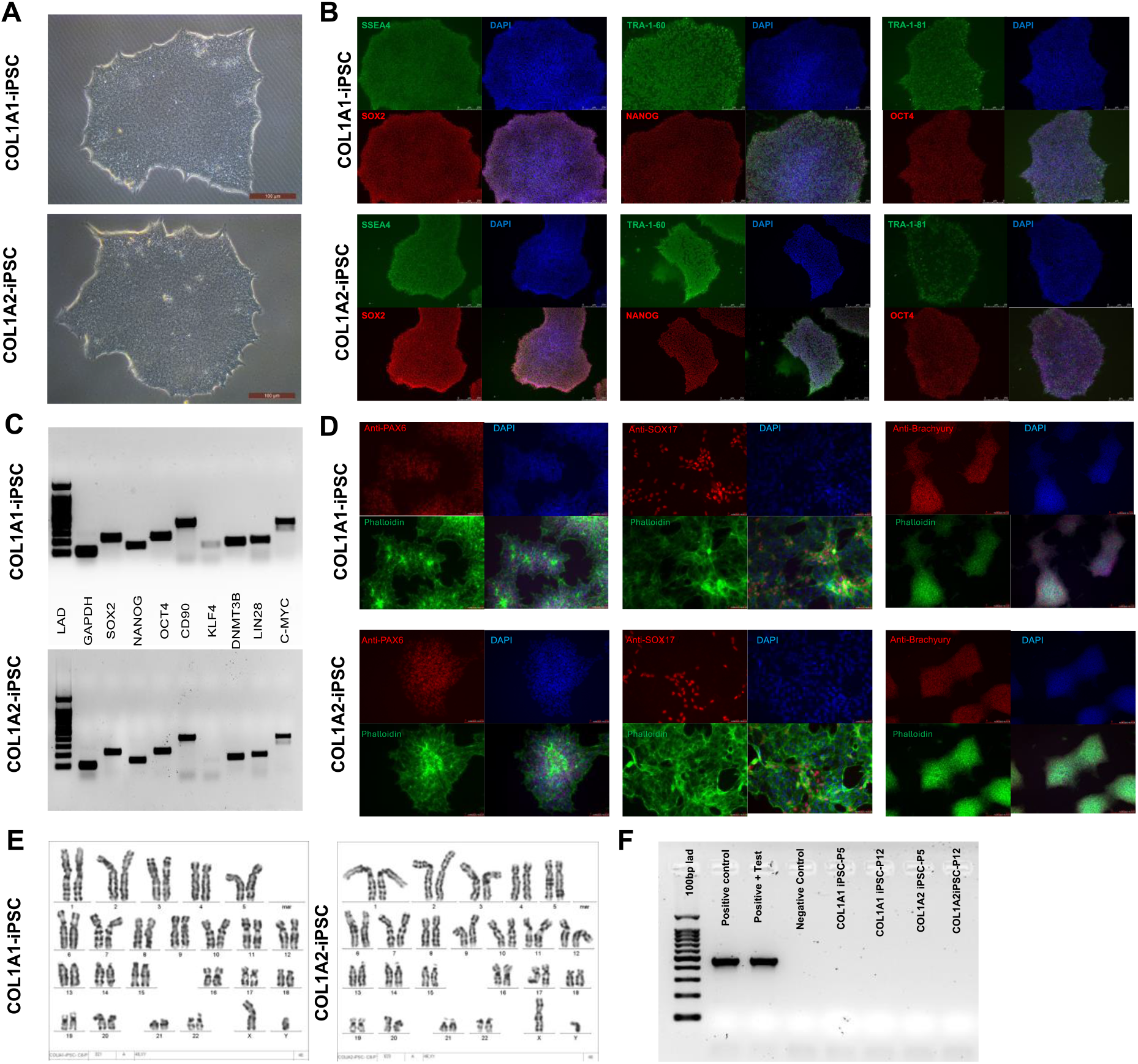
Characterization of OI Patient-Derived Induced Pluripotent Stem Cells (OI-iPSCs). (A) Phase-contrast microscopy image of OI-iPSC colonies (Passage 12) exhibiting typical pluripotent stem cell morphology, including distinct, smooth-edged colonies and a high nucleus-to-cytoplasm ratio. (B) Immunofluorescence staining demonstrating expression of the pluripotency markers SSEA4, TRA-1-60, and TRA-1-81 (green), along with the nuclear transcription factors SOX2, OCT4, and NANOG (red), in both OI-iPSC lines (COL1A1 and COL1A2) at Passage 12. (C) Semiquantitative RT-PCR analysis confirming expression of key pluripotency-associated genes (SOX2, NANOG, OCT4, KLF4, DNMT3B, LIN28) in both OI-iPSC lines at Passage 12. (D) Immunofluorescence staining confirms the potential to differentiate into all three germ layers, indicated by expression of nuclear marker PAX6 (ectoderm), Brachyury (mesoderm), and SOX17 (endoderm) (red). (E) Karyotype analysis reveals normal chromosomes in both OI-iPSC lines. (F) Routine testing confirmed the absence of mycoplasma contamination.

### iMSC derived osteogenic cells recapitulate the OI disease phenotype

At first, we attempted direct differentiation of OI patient-derived iPSCs into osteoblasts following a published protocol^16^. COL1A1 and COL1A2-OI iPSCs differentiated and formed mineralized monolayers of osteoblasts by day 10, as indicated by the Alizarin Red staining. However, compared to a control hiPSC, a significant decrease in the rate of mineralization was observed (supplementary fig. 2). However, the direct differentiation of iPSCs to osteoblasts using retinoic acid failed to recapitulate the disease phenotype observed in primary BMMSCs from OI patients (Figure 1). The various differentiation protocols for iPSCs to osteoblasts have significant limitations; they yield cells with varying osteogenic potential and may not consistently produce optimal osteoblasts. The variability in the differentiation methods also implies challenges in achieving reliable and effective osteoblast generation from iPSCs ^9^. Given the limitations of direct differentiation of iPSCs to osteoblasts, OI-iPSCs were differentiated into induced mesenchymal stem cells (iMSCs) through a two-step process. First, single-cell suspensions of OI-iPSCs were differentiated into early mesodermal progenitor cells. Second, these early mesodermal progenitor cells were further differentiated into early mesenchymal progenitor cells. The resulting cells were then expanded through multiple passages to obtain a homogeneous population of induced mesenchymal stromal cells (iMSCs). A typical spindle-shaped fibroblast-like morphology was observed under a phase-contrast microscope (Fig. 3A). Both sources of OI-iMSCs demonstrated the ability to differentiate into chondrocytes and adipocytes, as evidenced by the presence of glycosaminoglycans in Safranin O staining and lipid vacuoles in Oil Red O staining. Similar to OI-BMMSC, impaired osteogenic differentiation and poor mineralization were observed in COL1A1 and COL1A2 OI-iMSCs (Fig 3B). Flow cytometric analysis of OI-iMSCs revealed more than 98% expression of MSC surface markers CD73, CD90, CD105, and CD44 and less than 1% expression of pluripotent markers and negative markers TRA-1-60, CD24, CD117, CD45, CD34, and HLA-DR (Fig 3C). Gene expression analysis of chondrogenic, adipogenic, and osteogenic genes showed downregulation of SOX2, ACAN, PPARy, FABP4, RUNX2, ALP, and OCN compared to healthy BMMSC (p < 0.05, p < 0.005, p < 0.001, p < 0.0001) (Fig 3D). Sanger’s sequencing of the COL1A1 and COL1A2 loci in iMSC confirmed the integrity of the patient cell-specific mutations (Fig 3E,F).

**Figure 3.**
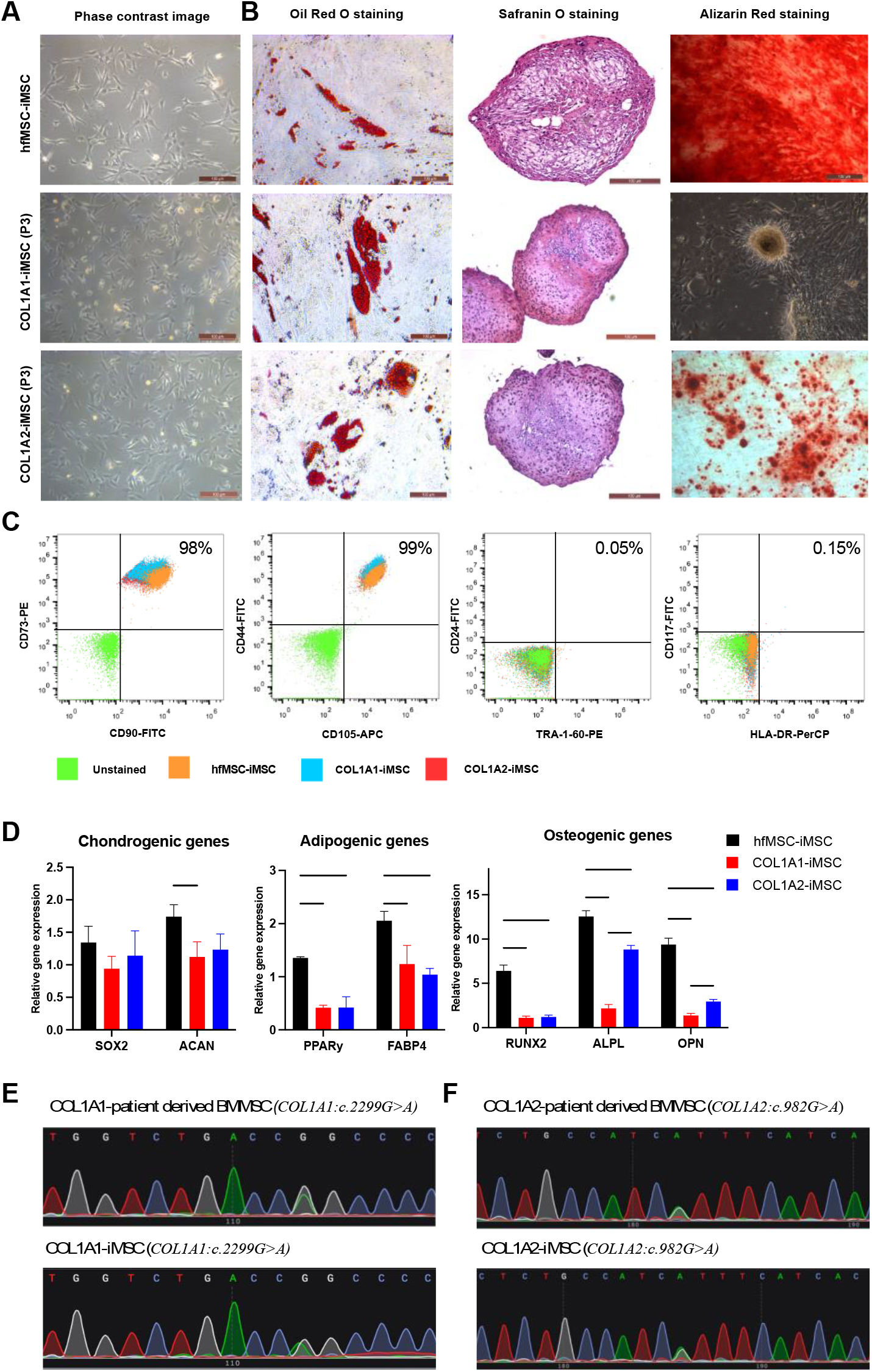
Characterization of OI Patient-Derived Induced Mesenchymal Stem Cells (OI-iMSCs). (A) Phase-contrast microscopy images showing the typical spindle-shaped morphology of both COL1A1 and COL1A2 OI-iMSC lines at Passage 3, comparable to healthy BMMSCs. (B) Trilineage differentiation assays demonstrating adipogenesis (Oil Red O staining: lipid vacuoles) and chondrogenesis (Safranin O staining: glycosaminoglycans). Osteogenesis (Alizarin Red staining) showed robust calcium deposition in healthy control BMMSCs but impaired mineralization in OI-iMSCs. (C) Flow cytometric analysis confirming high expression of positive MSC surface markers (CD90, CD105, CD73, CD44) and lack of expression of pluripotency markers (CD24, CD117, and TRA-1-60) in OI-iMSCs. (D) Gene expression analysis following differentiation. Key chondrogenic (SOX2, ACAN) and adipogenic (PPARγ, FABP4) genes were expressed. In contrast, osteogenic genes (RUNX2, ALPL, OPN) were significantly downregulated in OI-iMSCs compared to healthy controls at Day 21 of osteogenic induction (p < 0.05, *p < 0.001, ***p < 0.0001). (E, F) Sanger sequencing chromatograms confirming the presence of the patient-specific missense mutations in the COL1A1-OI-iMSC line (E) and the COL1A2-OI-iMSC line (F).

### Functional characterization of osteogenic cells derived from OI-iMSCs

To assess the disease phenotype, OI-iMSCs were subjected to osteogenic differentiation and compared to healthy BMMSCs. The osteogenic differentiation results were similar to those observed in patient-derived BMMSCs. Specifically, COL1A1-iMSCs exhibited impaired osteogenic differentiation, characterized by increased cell death (10-15%) and rolling of cells from day 5 of differentiation (Fig 4A). COL1A2-iMSCs differentiated into osteoblasts but displayed poor calcium deposition, as indicated by Alizarin Red S staining (Fig 4A). Quantification of alizarin red staining at day 7, day 14, and day 21 showed a significant decrease in the rate of mineralization compared to the healthy BMMSCs (p<0.0001) (Fig 4B). Gene expression analysis of key osteogenic factors showed abnormal altered expression of early (RUNX2, ALP, COL1) and mineralization (OSX, OPN) genes (p<0.05, p<0.0001) (Fig 4C). The COL1A1 OI-iMSC demonstrated an abnormal early expression of the mineralization gene OPN. A 2-fold increase in OPN expression at day 7 and a 2-fold decrease at day 21 were observed in COL1A1 OI-iMSCs compared to healthy BMMSCs (Fig 4C). These results demonstrated the ability of iMSCs to recapitulate the disease phenotype observed in patient-derived BMMSCs.

**Figure 4.**
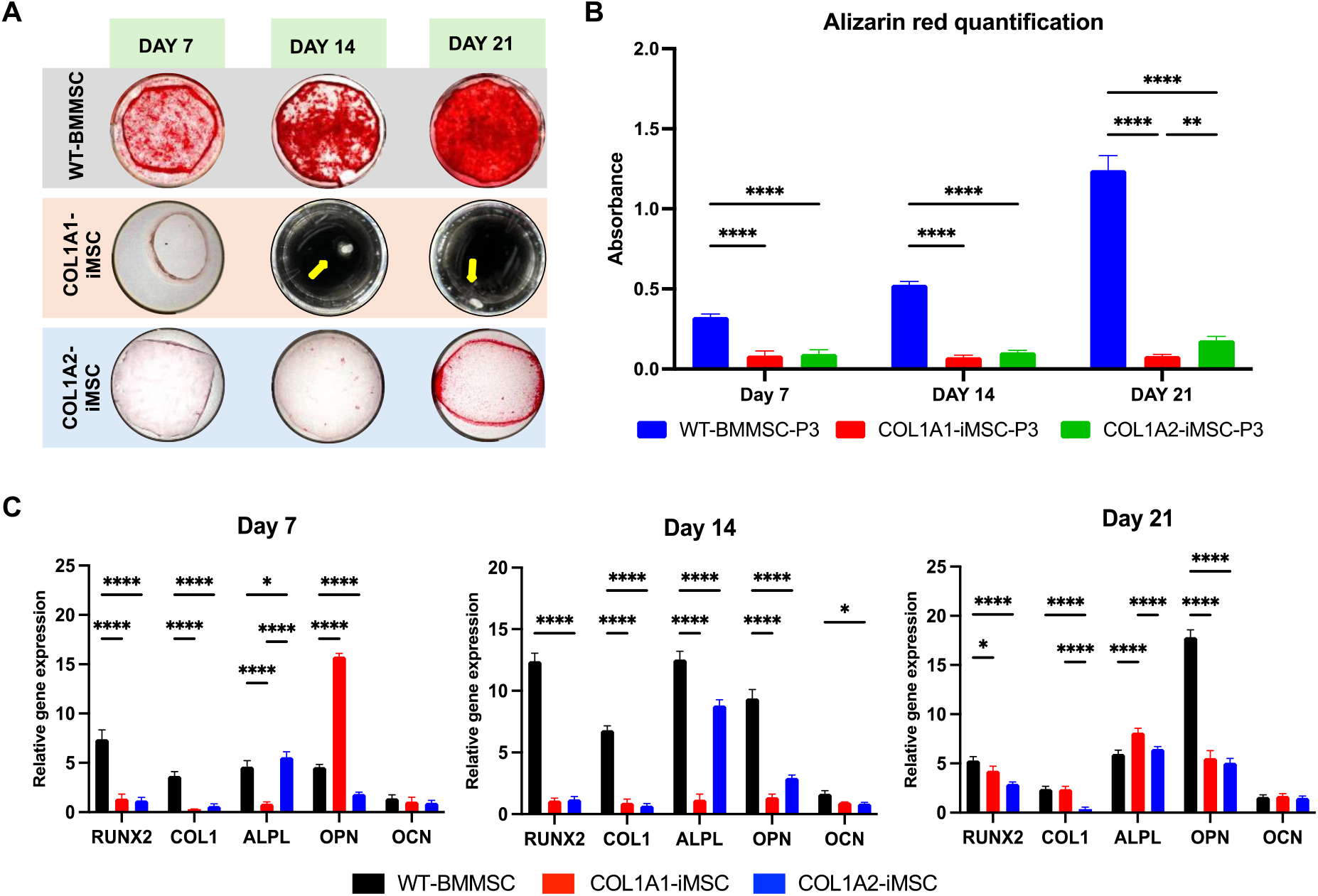
Functional Validation of the OI-iMSC Model during Osteogenic Differentiation. (A) Representative images of Alizarin Red staining during osteogenic differentiation of OI-iMSCs and healthy control BMMSCs at days 7, 14, and 21. Note the cellular detachment and rolling observed in COL1A1-iMSCs starting from day 7. COL1A2-iMSCs exhibit progressively impaired mineralization over 21 days compared to the robust mineralization in healthy controls. (B) Quantification of Alizarin Red staining revealing a significant decrease in calcium deposition in both COL1A1 and COL1A2-iMSC lines compared to controls at specified time points (**p < 0.01, ****p < 0.0001). (C) Gene expression analysis of key osteogenic markers by RT-qPCR. Expression levels of RUNX2, COL1, ALPL, and OCN were significantly decreased in OI-iMSCs compared to healthy controls at days 14 and 21. Notably, an aberrant early increase in OPN expression was observed specifically in COL1A1-iMSCs at day 7, which decreased at day 21. (*p<0.05, ****p<0.0001).

## Discussion

Primary MSCs have several limitations that restrict their use in understanding the disease mechanism of rare genetic bone disorders. These limitations include heterogeneous population, donor variability, impaired differentiation potential upon expansion, and lack of proliferation, as well as challenges related to the availability of donors, isolation, and expansion ^6,17^ Additionally, significant donor-specific variation can affect the difference in outcome analysis and reproducibility. Since the advent of induced pluripotent stem cell technology, a substantial number of patient-specific iPSC lines have been created for a wide range of rare genetic disorders. However, establishing reliable human disease models using these cells has been impeded by difficulties in controlling cell differentiation and determining disease-related mechanisms ^18,19^. Our study demonstrates the successful generation and application of patient-derived induced mesenchymal stem cells (iMSCs) as an in vitro model for Osteogenesis Imperfecta (OI), providing a robust platform for disease modeling and therapeutic development (Fig 5). The reprogramming of patient mesenchymal stromal cells into induced pluripotent stem cells (iPSCs) followed by differentiation into iMSCs through the mesoderm lineage ensures high fidelity to the patient’s genetic background and enables the study of OI-specific cellular phenotypes without losing the molecular and cellular phenotype ^20^. OI-iMSCs exhibit hallmark disease characteristics, including impaired osteogenic differentiation and reduced mineralization, which are consistent with the clinical manifestations of OI. Using iPSC-derived iMSC, the limitations of using the primary BMMSCs, i.e., heterogeneous population, donor variability, impaired differentiation potential upon expansion, and lack of proliferation, are circumvented ^21^.

**Figure 5.**
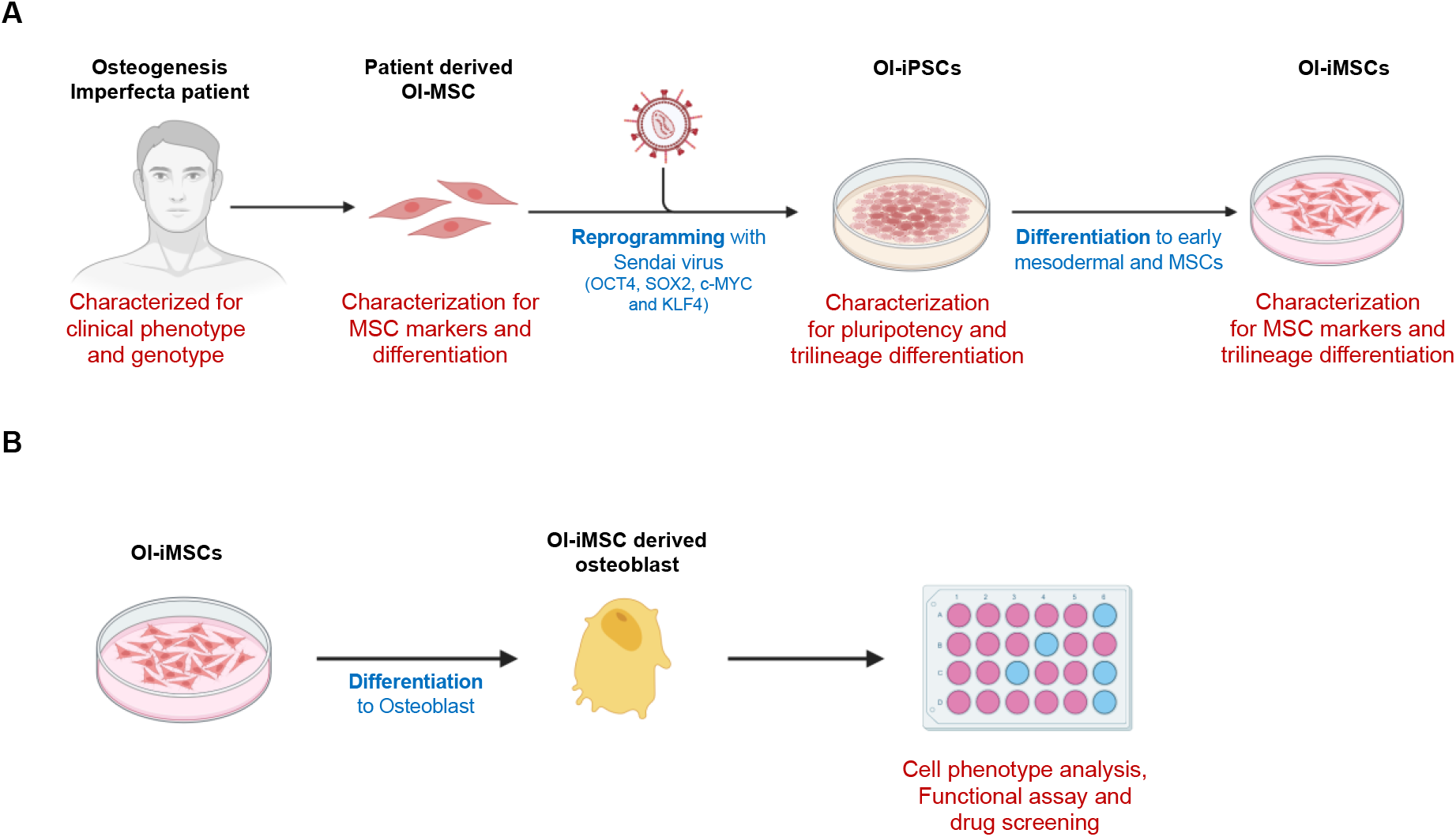
Overview of Experimental Workflow. (A) Patients with clinical presentations of Osteogenesis Imperfecta (OI) were selected based on clinical assessment and genotyping. Bone marrow mesenchymal stromal cells (BMMSCs) were isolated from OI patients harboring COL1A1 or COL1A2 variants and subsequently characterized. Using Sendai virus transduction, OI-BMMSCs were reprogrammed into induced pluripotent stem cells (OI-iPSCs), which were then characterized for pluripotency and trilineage differentiation potential. Finally, OI-iPSCs were differentiated into induced mesenchymal stem cells (OI-iMSCs) via a two-step protocol, followed by characterization for MSC surface marker expression and trilineage differentiation capacity, and validation for the OI disease phenotype.

We established two different iPSC-based cellular disease models of osteogenesis imperfecta with missense variants in the COL1A1:p.(Gly767Ser) and COL1A2:p.(Gly328Ser) genes. Most of the variants found in COL1A1/A2 are glycine substitutions, of which more than 95% of the patients present a lethal OI phenotype ^22^. These variants, affecting conserved glycine residues within the triple helical region of type I collagen, are predicted to cause substantial asymmetric disruptions to the triple helix structure ^23,24^. Previous experimental evidence indicates that glycine-to-serine mutations are frequently associated with a lethal OI phenotype ^22,25,26^ Both the iPSC disease models expressed pluripotent characteristics, including morphology, pluripotent markers, and trilineage differentiation into ectoderm, endoderm, and mesoderm lineages. The patient’s specific mutation was retained in the iPSC lines and the differentiated iMSCs with a normal karyotype. We differentiated the iPSCs via early mesodermal and early mesenchymal progenitors into mesenchymal stromal cells using animal component-free conditions. The differentiated iMSCs showed a homogenous population of distinct spindle-shaped fibroblast-like cells with adherence to the plastic surface, expression of MSC surface markers, and differentiation to chondrocytes, adipocytes, and osteoblasts with impaired mineralization. iMSCs offer a valuable source of patient-specific cells for developing effective treatments for osteogenesis imperfecta (OI) and other rare bone disorders, unlike fibroblast cell models, which fail to capture the complex pathophysiology of bone tissue.

We observed significant cell death in the COL1A1 and COL1A2 iMSCs starting from day 3 of osteogenic differentiation; 15-20% cell death was observed in COL1A1 and 5-10% in COL1A2. Our findings align with previous studies reporting apoptosis in other osteogenesis imperfecta models harboring mutations in the α-1 and α-2 collagen chains ^26–28^. The COL1A1-OI-iMSCs displayed impaired attachment to the culture surface starting from day 5 during the early osteoblast differentiation, leading to cellular rolling and pellet formation. While the COL1A2-OI-iMSCs were able to attach and grow as a monolayer for 21 days with impaired mineralization. Gene expression analysis showed significant downregulation of osteogenic markers COL1, RUNX2, ALP, OCN, and OPN in our cellular disease model, as reported in [30]. The decreased expression of COL1A1/A2 results from the downregulation of the key osteogenic transcription factor RUNX2. Alizarin red staining demonstrated reduced calcium mineralization during bone formation, consistent with the low bone mass and density seen in OI. This impaired mineralization may underlie the observed decreases in bone cells, collagen production, poor extracellular matrix mineralization, and increased cell death due to ER stress. Our data provides clear evidence that the pathogenesis associated with COL1A1:p.(Gly767Ser) and COL1A2:p.(Gly328Ser) can be modeled in the osteoblast lineages generated from the iMSCs derived from the patients with type III and type IV osteogenesis imperfecta.

Previous studies on disease modeling of OI using iPSCs revealed mineralization defects and impaired collagen synthesis ^8,16^. In fact, in our study when the OI-iPSCs were directly differentiated to osteogenic cells, the results contradicted our observations from BMMSC differentiation, with mineralization defects observed in both COL1A1 and COL1A2. However, the two-stage differentiation of OI-iPSC to iMSC and to osteoblast demonstrated impaired mineralization and recapitulated the disease phenotype seen in the OI-patient-derived BMMSCs. The iMSC-based approach is more reliable, cost-effective, and provides a homogenous, scalable cell population, particularly suited for modeling musculoskeletal diseases such as osteogenesis imperfecta. Also, iMSC offers a robust platform for high-throughput screening of drug molecules ^20,29^.

## Conclusion

Our study has successfully established two novel iPSC-based cellular disease models for osteogenesis imperfecta, accurately replicating the disease phenotype. Creating patient-specific iMSCs offers a minimally invasive approach to studying osteogenesis imperfecta (OI). Unlike primary cell culture models, iMSCs display better potency allowing efficient upscaling of cells if required. This provides an opportunity for personalized investigation of osteoblasts, collagen production, and drug screening, potentially accelerating the development of targeted OI therapies.

---

**Supplementary figure 1:**
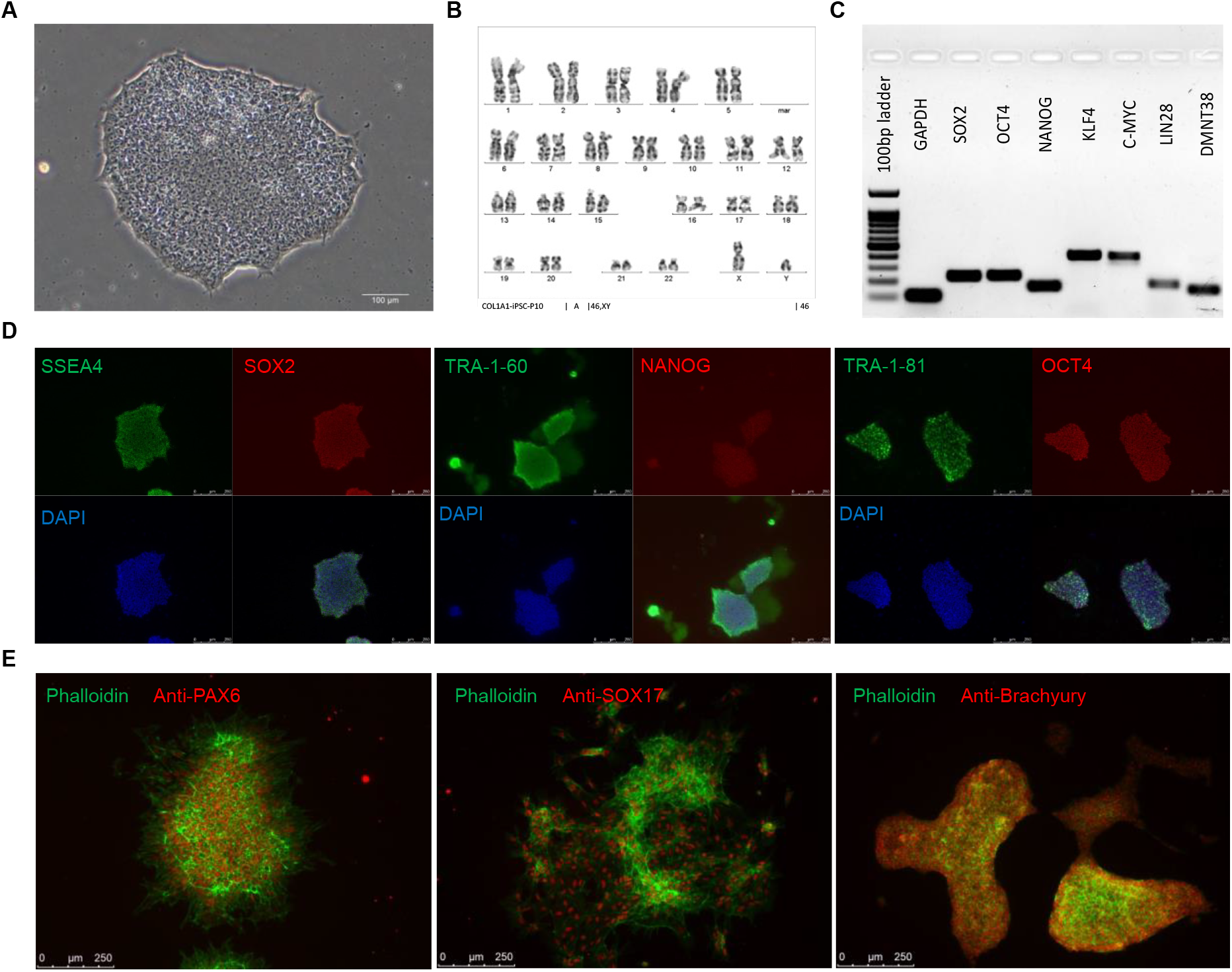
Characterization of hfMSC derived iPSC. (A) Phase-contrast microscopy image of hfMSC-iPSC colonies (Passage 10) exhibiting typical pluripotent stem cell morphology with distinct smooth-edged colonies and a high nucleus-to-cytoplasm ratio. (B) Karyotype analysis reveals a normal chromosome. (C) Semiquantitative RT-PCR analysis confirms expression of key pluripotency-associated genes (SOX2, NANOG, OCT4, KLF4, DNMT3B, LIN28) in both hfMSC-iPSC at Passage 10. (D) Immunofluorescence staining demonstrates the expression of pluripotency markers SSEA4, TRA-1-60, and TRA-1-81 (green), along with the nuclear transcription factors SOX2, OCT4, and NANOG (red) at passage 10. (D) Immunofluorescence staining following in trilineage differentiation confirms the potential to differentiate into all three germ layers, indicated by expression of nuclear marker PAX6 (ectoderm), SOX17 (endoderm), and Brachyury (mesoderm).

**Supplementary figure 2:**
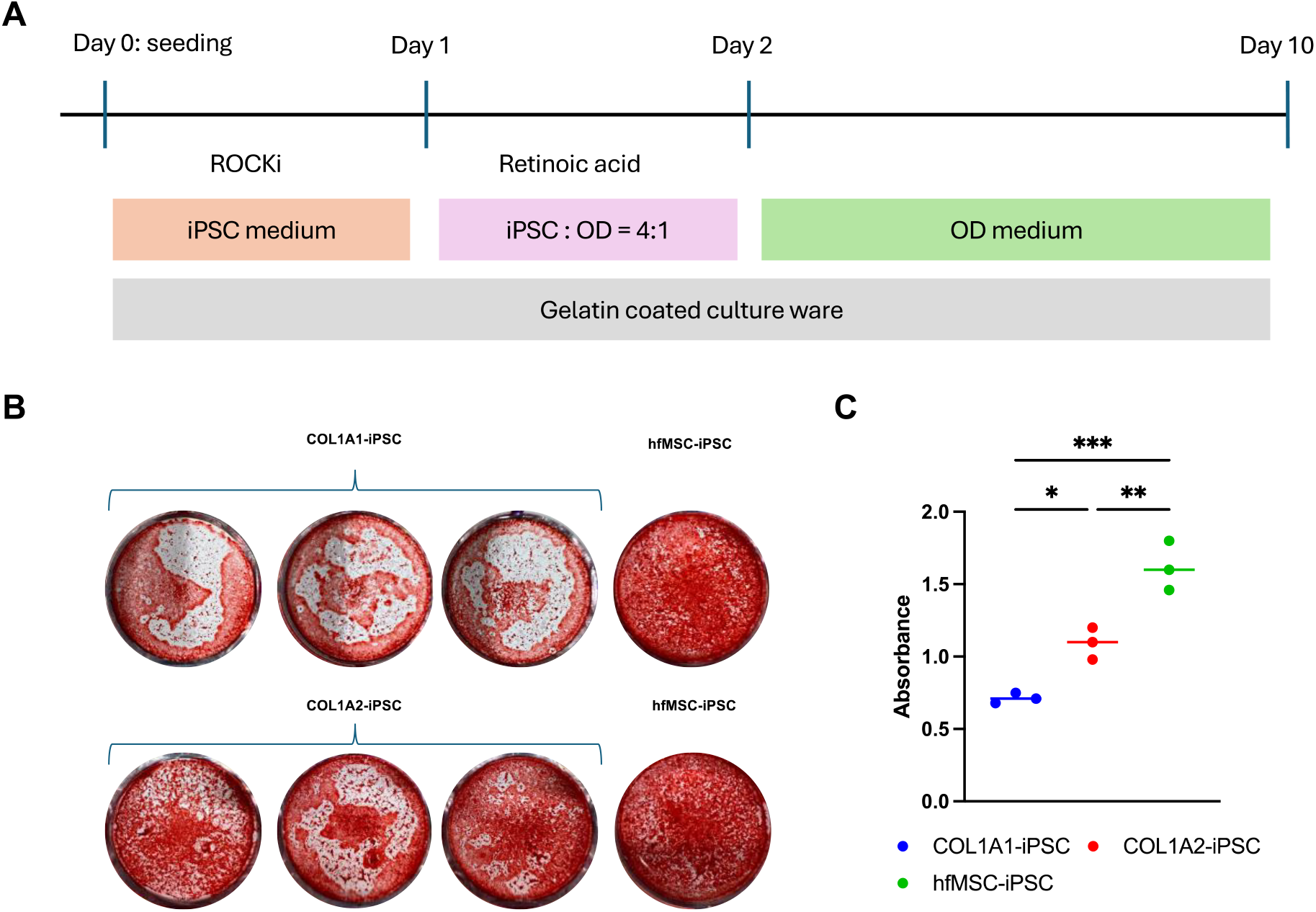
Direct differentiation of iPSC to osteoblast. A) Schematic illustration of differentiation protocol. B) Alizarin red staining at day 10 of osteogenic differentiation of COL1A1-iPSCs, COL1A2-iPSCs, and control hfMSC derived iPSCs shows calcium nodule formation. C) Quantification of alizarin red staining shows a decrease in the rate of mineralization in OI-iPSCs compared to the control hfMSC-iPSCs. (*p<0.05, **p<0.01, ***p<0.001)

## References

1. Forlino A, Marini JC, Branch EM, Shriver EK. Osteogenesis imperfecta. Lancet. 2015;387(10028):1657. doi:10.1016/S0140-6736(15)00728-X

2. Marini JC, Forlino A, Bächinger HP, et al. Osteogenesis imperfecta. Nat Rev Dis Primers. 2017;3. doi:10.1038/NRDP.2017.52

3. Marie PJ. Osteoblast dysfunctions in bone diseases: from cellular and molecular mechanisms to therapeutic strategies. Cell Mol Life Sci. 2015;72(7):1347–1361. doi:10.1007/S00018-014-1801-2

4. Schott T, Eisenberg KA, Vuillermin CB, Bae DS, Waters PM, Bauer AS. Donor-Site Morbidity for Iliac Crest Harvesting for Pediatric Scaphoid Nonunion. J Hand Surg Am. 2023;48(8):833.e1–833.e5. doi:10.1016/J.JHSA.2022.02.007

5. Shin SR, Tornetta P. Donor Site Morbidity After Anterior Iliac Bone Graft Harvesting. J Orthop Trauma. 2016;30(6):340–343. doi:10.1097/BOT.0000000000000551

6. Wagner W, Horn P, Castoldi M, et al. Replicative senescence of mesenchymal stem cells: A continuous and organized process. PLoS One. 2008;3(5). doi:10.1371/JOURNAL.PONE.0002213

7. Q L, Y Z, X L, F G, HF T. Directed Differentiation of Human-Induced Pluripotent Stem Cells to Mesenchymal Stem Cells. Methods Mol Biol. 2016;1416:289–298. doi:10.1007/978-1-4939-3584-0_17

8. Takeyari S, Kubota T, Ohata Y, et al. 4-Phenylbutyric acid enhances the mineralization of osteogenesis imperfecta iPSC-derived osteoblasts. J Biol Chem. 2020;296:100027. doi:10.1074/JBC.RA120.014709

9. Wu Q, Yang B, Hu K, Cao C, Man Y, Wang P. Deriving Osteogenic Cells from Induced Pluripotent Stem Cells for Bone Tissue Engineering. https://home.liebertpub.com/teb. 2017;23(1):p1-8. doi:10.1089/TEN.TEB.2015.0559

10. Duangchan T, Tawonsawatruk T, Angsanuntsukh C, et al. Amelioration of osteogenesis in iPSC-derived mesenchymal stem cells from osteogenesis imperfecta patients by endoplasmic reticulum stress inhibitor. Life Sci. 2021;278. doi:10.1016/J.LFS.2021.119628

11. Claeys L, Zhytnik L, Ventura L, et al. In Vitro Modelling of Osteogenesis Imperfecta with Patient-Derived Induced Mesenchymal Stem Cells. Int J Mol Sci. 2024;25(6). doi:10.3390/ijms25063417

12. Dias IX, Cordeiro A, Guimarães JAM, Silva KR. Potential and Limitations of Induced Pluripotent Stem Cells-Derived Mesenchymal Stem Cells in Musculoskeletal Disorders Treatment. Biomolecules. 2023;13(9):1342. doi:10.3390/BIOM13091342

13. Li C, Zhao H, Wang B. Mesenchymal stem/stromal cells: Developmental origin, tumorigenesis and translational cancer therapeutics. Transl Oncol. 2021;14(1):100948. doi:10.1016/J.TRANON.2020.100948

14. Pierini M, Dozza B, Lucarelli E, et al. Efficient isolation and enrichment of mesenchymal stem cells from bone marrow. Cytotherapy. 2012;14(6):686–693. doi:10.3109/14653249.2012.677821

15. Frobel J, Hemeda H, Lenz M, et al. Epigenetic rejuvenation of mesenchymal stromal cells derived from induced pluripotent stem cells. Stem Cell Reports. 2014;3(3):414–422. doi:10.1016/J.STEMCR.2014.07.003

16. Kawai S, Yoshitomi H, Sunaga J, et al. In vitro bone-like nodules generated from patient-derived iPSCs recapitulate pathological bone phenotypes. Nat Biomed Eng. 2019;3(7):558–570. doi:10.1038/S41551-019-0410-7

17. Wagner W, Feldmann RE, Seckinger A, et al. The heterogeneity of human mesenchymal stem cell preparations—Evidence from simultaneous analysis of proteomes and transcriptomes. Exp Hematol. 2006;34(4):536–548. doi:10.1016/J.EXPHEM.2006.01.002

18. Tang S, Xie M, Cao N, Ding S. Patient-Specific Induced Pluripotent Stem Cells for Disease Modeling and Phenotypic Drug Discovery. J Med Chem. 2016;59(1):2–15. doi:10.1021/ACS.JMEDCHEM.5B00789/ASSET/IMAGES/MEDIUM/JM-2015-00789Q_0002.GIF

19. Grskovic M, Javaherian A, Strulovici B, Daley GQ. Induced pluripotent stem cells--opportunities for disease modelling and drug discovery. Nat Rev Drug Discov. 2011;10(12):915–929. doi:10.1038/NRD3577

20. Zhang J, Chen M, Liao J, et al. Induced Pluripotent Stem Cell-Derived Mesenchymal Stem Cells Hold Lower Heterogeneity and Great Promise in Biological Research and Clinical Applications. Front Cell Dev Biol. 2021;9. doi:10.3389/FCELL.2021.716907

21. Wruck W, Graffmann N, Spitzhorn LS, Adjaye J. Human Induced Pluripotent Stem Cell-Derived Mesenchymal Stem Cells Acquire Rejuvenation and Reduced Heterogeneity. Front Cell Dev Biol. 2021;9. doi:10.3389/FCELL.2021.717772

22. Garibaldi N, Besio R, Dalgleish R, et al. Dissecting the phenotypic variability of osteogenesis imperfecta. Dis Model Mech. 2022;15(5):dmm049398. doi:10.1242/DMM.049398

23. Bodian DL, Madhan B, Brodsky B, Klein TE. Predicting the clinical lethality of osteogenesis imperfecta from collagen glycine mutations. Biochemistry. 2008;47(19):5424–5432. doi:10.1021/BI800026K/SUPPL_FILE/BI800026K-FILE002.PDF

24. Glycine to serine substitution in the triple helical domain of pro-alpha 1 (II) collagen results in a lethal perinatal form of short-limbed dwarfism - PubMed. Accessed March 21, 2025. https://pubmed.ncbi.nlm.nih.gov/2572591/#

25. Pack M, Constantinou CD, Kalia K, Nielsen KB, Prockop DJ. Substitution of serine for α1(I)-glycine 844 in a severe variant of osteogenesis imperfecta minimally destabilizes the triple helix of type I procollagen. The effects of glycine substitutions on thermal stability are either position or amino acid specific. Journal of Biological Chemistry. 1989;264(33):19694–19699.

26. Westerhausen A, Kishi J, Prockop DJ. Mutations that substitute serine for glycine α1-598 and glycine α-631 in type I procollagen: The effects on thermal unfolding of the triple helix are position-specific and demonstrate that the protein unfolds through a series of cooperative blocks. Journal of Biological Chemistry. 1990;265(23):13995–14000. doi:10.1016/s0021-9258(18)77447-4

27. Gioia R, Panaroni C, Besio R, et al. Impaired Osteoblastogenesis in a Murine Model of Dominant Osteogenesis Imperfecta: A New Target for Osteogenesis Imperfecta Pharmacological Therapy. Stem Cells. 2012;30(7):1465–1476. doi:10.1002/STEM.1107

28. Lisse TS, Thiele F, Fuchs H, et al. ER stress-mediated apoptosis in a new mouse model of osteogenesis imperfecta. PLoS Genet. 2008;4(2). doi:10.1371/JOURNAL.PGEN.0040007

29. Lo Cicero A, Jaskowiak AL, Egesipe AL, et al. A High Throughput Phenotypic Screening reveals compounds that counteract premature osteogenic differentiation of HGPS iPS-derived mesenchymal stem cells. Sci Rep. 2016;6:34798. doi:10.1038/SREP34798

